# Targeting Vaccinations for the Licensed Dengue Vaccine: Considerations for Serosurvey Design

**DOI:** 10.1101/171140

**Authors:** N Imai, NM Ferguson

## Abstract

**Objective:** WHO recommends countries consider dengue vaccination in geographic settings only where epidemiological data indicate a high burden of disease. In defining target populations, WHO recommend that prior infection with any dengue serotype should be >70% seroprevalence. Here we address considerations for serosurvey design in the context of the newly licensed CYD-TDV vaccine.

**Methods:** To explore how the design of seroprevalence surveys (age range, survey size) would affect estimates of the force of infection, for every combination of age range, total survey size, transmission setting, and test sensitivity/specificity, 100 age-specific seroprevalence surveys were simulated using a beta-binomial distribution and a simple catalytic model. The transmission intensity was then re-estimated using a Metropolis-Hastings Markov Chain Monte-Carlo algorithm.

**Findings:** Sampling from a wide age range led to more accurate estimates than having a larger sample size. This finding was consistent across all transmission settings. The optimal age range to sample from differed by transmission intensity, with younger and older ages being important in high and low transmission settings respectively. The optimum test sensitivity and specificity given an imperfect test also differed by transmission setting with high sensitivity being important in high transmission settings and high specificity important in low transmission settings.

**Conclusions:** When assessing the suitability for vaccination by seroprevalence surveys, countries should ensure that an appropriate age range is sampled, taking into account epidemiological evidence about the local burden of dengue.

## Introduction

The total incidence of dengue cases has increased 30-fold in the past 50 years. Dengue infection is the cause of an estimated 1.14 million (0.73 million – 1.98 million) disability-adjusted life-years (DALYs) in 2013 [1]. Consequently, dengue is now the most important mosquito-borne viral infection globally, and with 40% of the world’s population at risk of infection, dengue imposes a significant public health burden [2,3].

The first dengue vaccine - the Sanofi CYD-TDV vaccine, has now been licensed by several dengue-endemic countries for use in 9-45 years and 9-60 year olds; including in Brazil, El Salvador, Paraguay [4], Costa Rica [5], Mexico [6], and the Philippines [7]. Results from large-scale phase III trials conducted in 2-14 year olds in 5 countries in Asia showed a reduction in hospitalisations and 80% efficacy against severe dengue [8]. The CYD14/15 trial conducted in 9-16 year olds in 5 countries in Latin America showed a vaccine efficacy of 80.8% (95% CI: 70.1 – 87.7) against any hospitalised dengue infection [9]. However the vaccine efficacy varied by infecting serotype, age, and baseline serostatus. Additionally, in vaccinated children aged 2-5 years in Asia, there was a statistically significant increased risk of hospitalised dengue in the third year after vaccination (first dose) [8].

Due to the safety concerns and the limited protective effect of the vaccine in low transmission settings, the Strategic Advisory Group of Experts (SAGE) recommended the introduction of the vaccine only in high endemic regions (>70% seroprevalence in the target age group, 9 years and above). Vaccination was not recommended where seroprevalence is <50% in the target age group [10]. Given this safety signal, and in the context of the SAGE recommendations, accurate assessment of the seroprevalence of the target population is essential prior to large-scale implementation of the vaccine.

Age-specific seroprevalence surveys can show considerable fluctuations with age due to the epidemic nature of dengue transmission as well as sampling variability. Therefore, the observed seroprevalence in a single age group may not be representative of the true seroprevalence of the target age group. A more accurate measure of dengue endemicity is the transmission intensity or force of infection (*λ*) defined as the per capita rate that susceptible individuals acquire infection [11]. It is a parameter that captures the dynamics of dengue transmission, and since it takes into account the susceptibility of the population, it can be used to compare the rate of dengue transmission in different groups [12]. Therefore, it is an important measure when determining the impact of interventions such as vaccination. Dengue force of infection can be estimated from non-serotype and serotype-specific age-stratified seroprevalence data [13]. However, accurate estimation of *λ* requires seroprevalence data from a wide range of ages. Figure 1 demonstrates how a small difference in seroprevalence at a single age (which may be empirically statistically significant) does not necessarily equate to similar forces of infection. Therefore, in the context of the CYD-TDV vaccine it is important to sample from a wide age group to accurately assess vaccination suitability.

**Figure 1:**
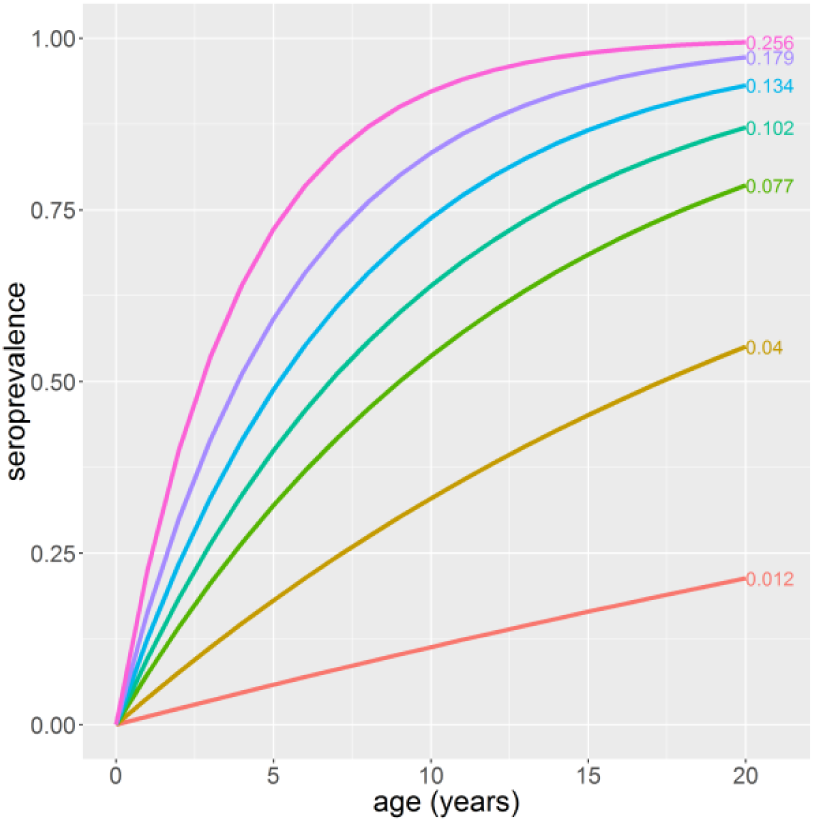
Change in seroprevalence with age at different transmission intensities. Transmission intensity values are given at the end of each seroprevalence curve.

In the context of the SAGE recommendations for the CYD-TDV vaccine, here we present considerations when designing seroprevalence surveys for targeting vaccination such as; the age-range from which we should sample, the survey size, and how the sensitivity and specificity of the assay may affect our estimates.

## Methods

### Model

Assuming that seroprevalence surveys are conducted using non-serotype specific assays such as the IgG enzyme-linked immunosorbent assays (ELISA), we assumed that the relationship between the seroprevalence and age is given by (1):

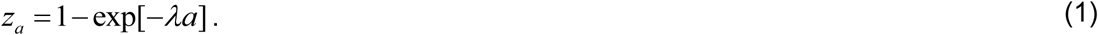

Where *z_a_* is the seroprevalence in age group *a*, *λ* is the force of infection or transmission intensity, and *a* is age in years. The model assumes a constant force of infection and generally describes the results of a cross-sectional IgG survey well [13].

### Simulation and Estimation Procedure

We considered 11 potential age ranges to test and 5 total survey sizes. We assumed that individuals were distributed uniformly across the age groups, i.e. for a total survey size of 2000 and an age range of 0 – 20 year olds; 96 individuals are tested at each age. We also considered 7 different transmission settings where the seroprevalence in 9 year olds (the target age group) ranged from 10% - 90%. The corresponding force of infection was calculated using equation 1. Finally, we looked at different test sensitivities and specificities (60% - 100%). Table 1 summarises the combinations tested.

**Table 1:**
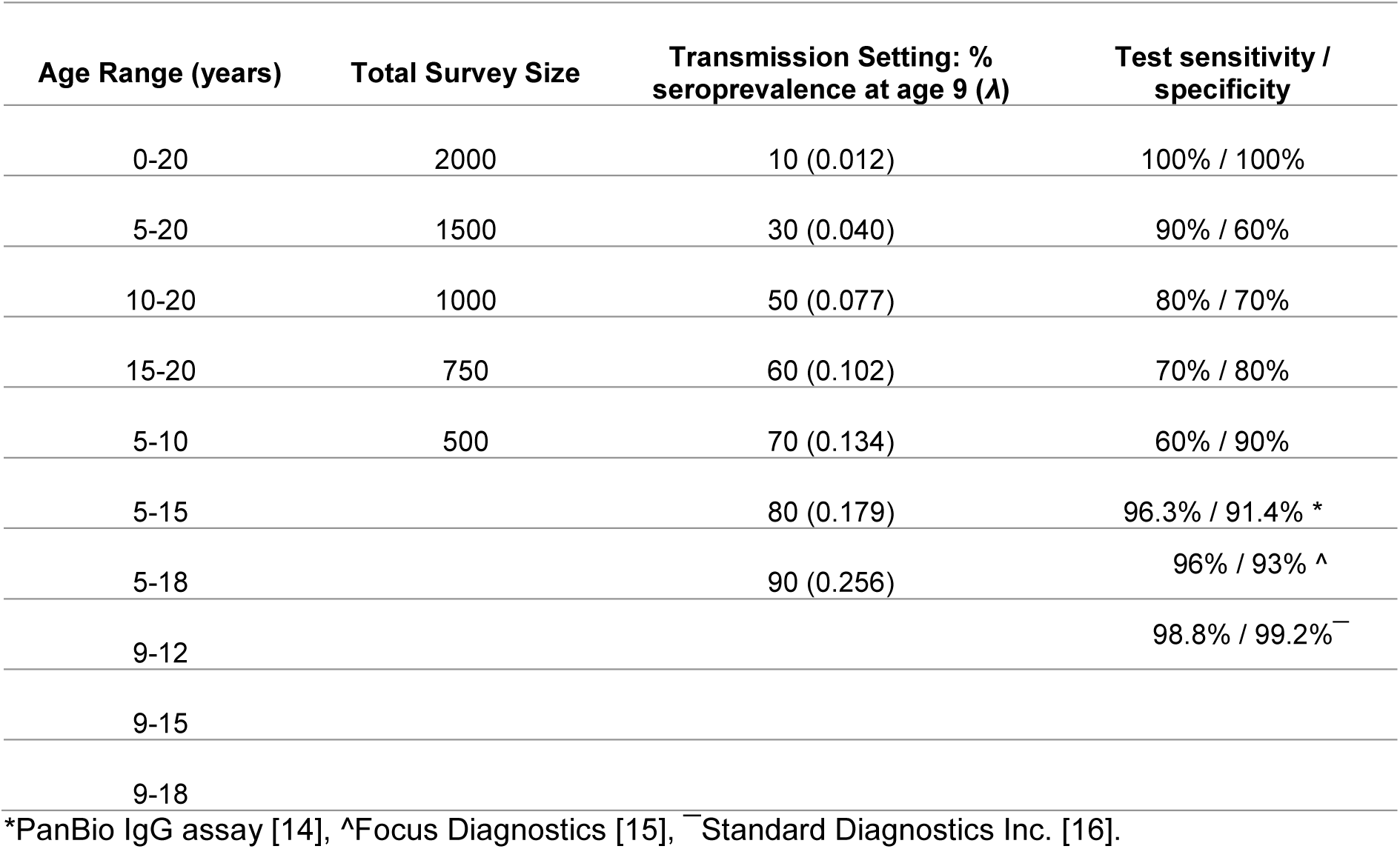
Summary of different scenarios used for simulated seroprevalence surveys

For every combination of age range, survey size, transmission setting, and test sensitivity-specificity, 100 seroprevalence surveys were simulated from a beta-binomial (BB) distribution which can account for overdispersion. We assumed that the probability of an individual in age group *a* being seropositive was beta-binomially distributed (2):

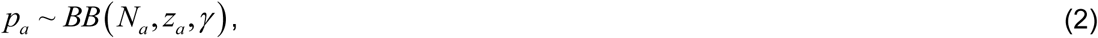

where *N_a_* is the number of individuals in age group *a*, *z_a_* is the proportion of the age group seropositive given by equation (1), and *γ* is over-dispersion (the smaller the value the larger the over-dispersion). Over-dispersion was fixed at *γ* = 25 (average value estimated from 28 seroprevalence surveys from 13 countries [13]).

The sensitivity and specificity will then affect whether an individual in age group *a* will test positive 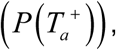 which is given by (3):

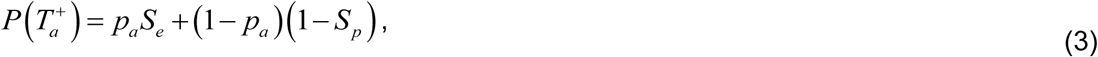

where *p_a_* is the probability an individual aged *a* is seropositive (2) and *S_e_* and *S_p_* are the sensitivity and specificity of the IgG test respectively. Thus, with a perfect test (100% sensitivity and specificity), 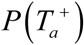 = *p_a_*. Figure 2 shows the probability of testing positive at different sensitivity and specificities given that the baseline probability of being seropositive is 0.7 (*p_a_* = 0.7) from a beta-binomial distribution. For comparison purposes, the same combinations were tested using a binomial distribution. For full details see SI text.

**Figure 2:**
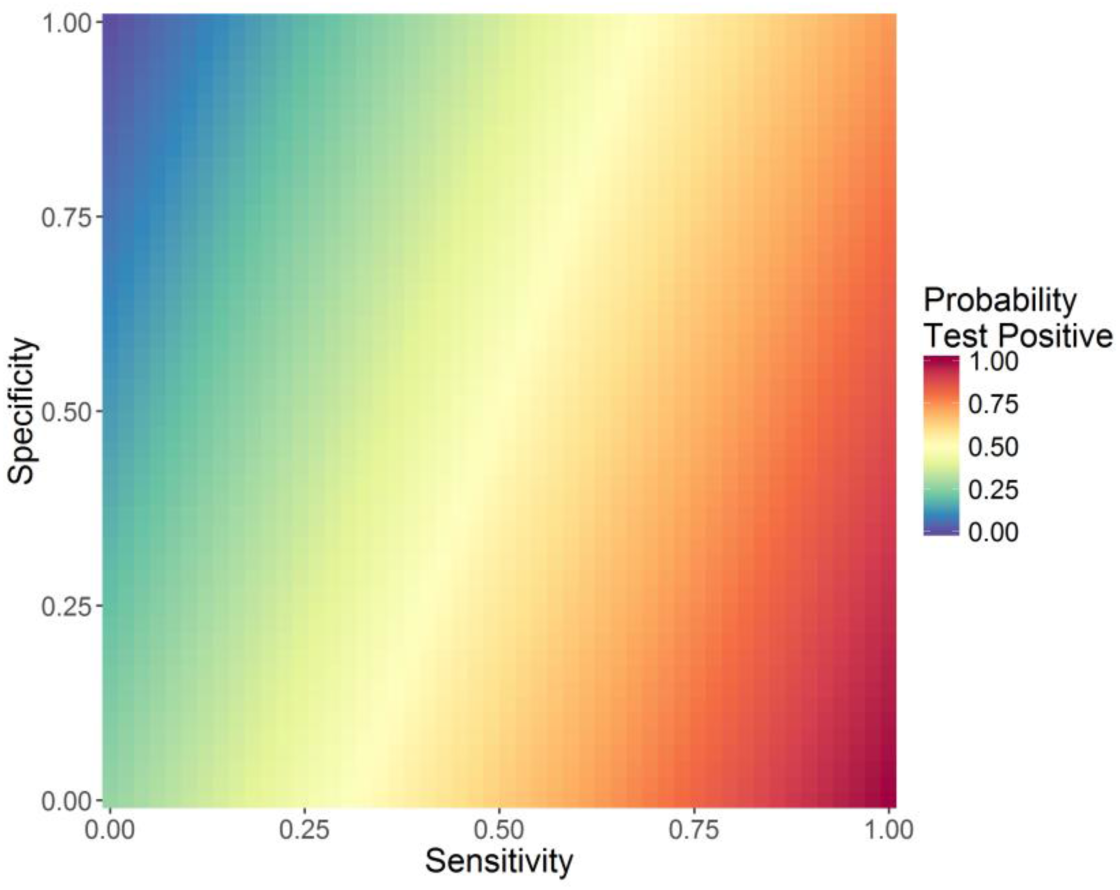
Probability of 9 year olds testing positive [P(T_a_^+^)] at different test sensitivity and specificities. Baseline probability of being seropositive is 0.7 from a beta-binomial distribution (p_9_ = 0.7).

The force of infection (*λ*) and overdispersion (*γ*) was then re-estimated from each simulated dataset (100 estimates per combination) using a Metropolis-Hasting Markov Chain Monte-Carlo (MH-MCMC) algorithm with a beta-binomial likelihood (4) and uniform priors.

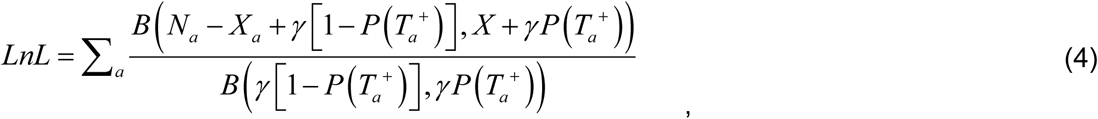

where *X_a_* is the number of individuals testing seropositive in age group *a*, *N_a_* is the total number of individuals tested in age group *a*, and 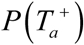 is the probability that an individual in age group *a* tests positive as defined above (3).

The model was fitted using the R Statistical Language (version 3.1.0) [17]. For details on the binomial estimation procedure see SI text.

The mean of the 100 mean posterior distributions (mean of the mean) and the standard deviation estimated from different combinations of age range, survey size, transmission setting, and test sensitivity and specificity were compared.

### Coefficient of Variation (relative standard deviation)

The average coefficient of variation (CV) gives a measure of variability in relation to the mean force of infection and is also a measure of the precision (the smaller the value, the more precise the estimate). The average CV was calculated over 100 simulated datasets for every combination of the force of infection, survey size, and sensitivity/specificity per age group.

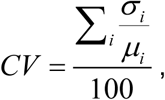

where *σ* is the standard deviation and *μ* is the mean.

## Results

With a high sensitivity and specificity test, across the different combinations of age-range, total survey size, and transmission setting, the true force of infection was correctly re-captured within the standard deviation of the 100 mean posterior distributions. However, the central estimate (mean of 100 means) was consistently overestimated when only sampling from a narrow age range e.g. 9-12 years or 15-20 years. As expected, the precision of the estimate increased with survey size for all transmission settings and age range sampled (smaller standard deviation around the central estimate) (Figure 3).

**Figure 3:**
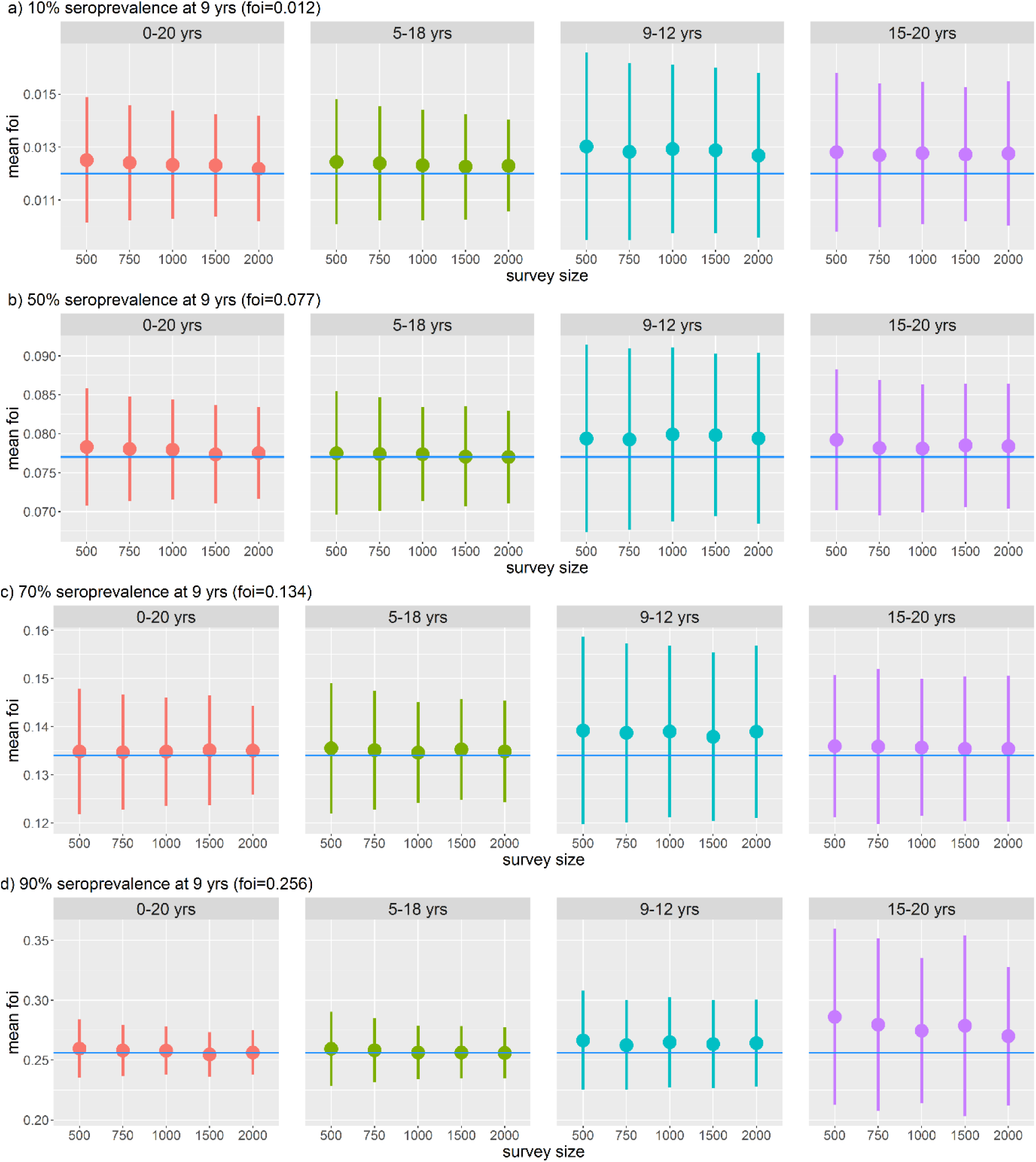
Dengue force of infection (*λ*) re-estimated from different age ranges and survey sizes at different transmission settings assuming a binomial distribution and a perfect test is used (100% sensitivity and specificity). A) Very low transmission setting (10% seroprevalence at 9 years), B) medium transmission setting (50% seroprevalence at 9 years), C) high transmission setting (70% seroprevalence at 9 years), and D) very high transmission setting (90% seroprevalence at 9 years). The point is the mean of the mean posterior estimates from 100 simulations and the line the standard deviation. The blue line shows the true value of *λ*.

In very low transmission settings (10% seroprevalence at 9 years), even with the widest age range (0-20 years) which included young children, the force of infection was overestimated (Figure 3a). Conversely in very high transmission settings (90% seroprevalence at 9 years), only sampling from older children 9 years and above overestimated the force of infection (Figure 3d).

Assuming seroprevalence surveys were carried out using a ‘perfect’ IgG assay (100% sensitivity, 100% specificity), the width of the age range sampled had a greater impact on the precision of the estimate compared to total sample size. With a narrow age range (e.g. 9-12 years), although the central estimate was close to the true value uncertainty around the estimate was large and could not be reduced regardless of survey size. This finding was consistent across all transmission settings (Figure 3a-d). This finding was not as marked when we did not account for overdispersion (SI text).

We also found that the optimum age-range to sample from differed by transmission setting. Figure 4 shows the average coefficient of variation (CV) for different age ranges and transmission settings. The coefficient of variation is a measure of the precision of our estimates where the smaller the value, the more precise the estimate. It represents the extent of variability in our force of infection estimates in relation to the mean. We found that in low transmission settings (FOI: 0.012, expected seroprevalence at age 9 = 10%) it is also important to sample from older age groups since the difference in seroprevalence amongst young children (<9 years) is small. Figure 4 shows that sampling 15-20 year olds gives a more precise estimate than sampling from 9-12 year olds in this setting. Conversely, in high transmission settings (FOI: 0.256, expected seroprevalence at age 9 = 90%) it is important to also sample from young age groups since differences in seroprevalence among older ages will be small. These results are also reflected in Figure 3a, where in very low transmission settings (10% seroprevalence at age 9) even with the widest age range 0-20 years the central estimate is overestimating the true force of infection. This suggests that at very low transmission settings the change in seroprevalence between 0 and 20 year olds is not sufficient to accurately estimate transmission intensity and older age groups (adults) need to be sampled.

**Figure 4:**
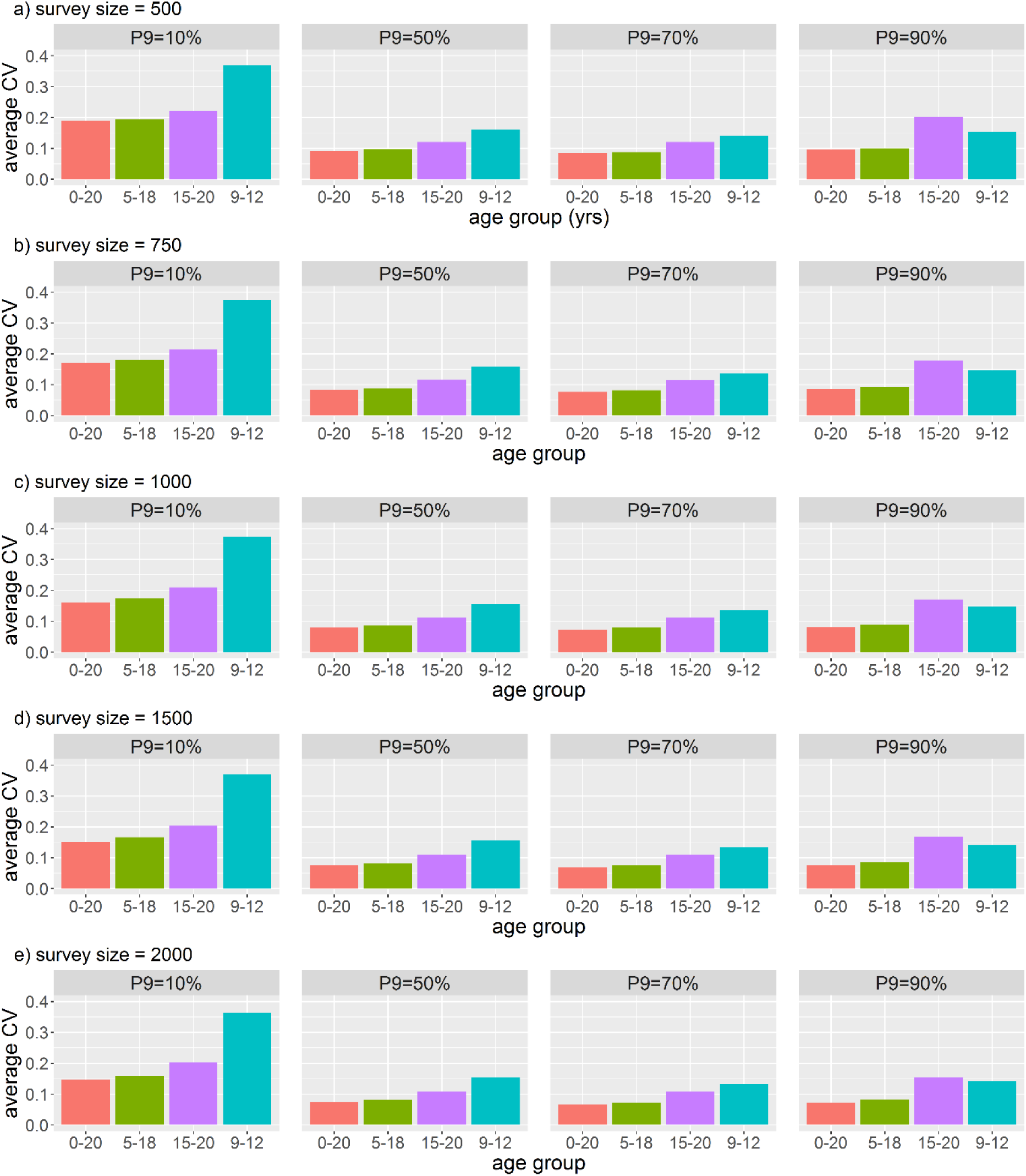
Average coefficient of variation (CV) for different age ranges sampled at different transmission settings (P9 = expected seroprevalence at age 9). Results assume a perfect test (100% sensitivity and specificity) is used. Panels show total survey sizes of a) 500; b) 750; c) 1000; d) 1500; and e) 2000.

We found that there was no substantial gain in precision of our estimates above a survey size of 1000. As expected, the larger the survey size, the smaller the CV or more precise our estimates.

Comparing the effect of different test sensitivities and specificities on the ability to accurately re-estimate the force of infection, as expected the perfect test (100% sensitivity and specificity) performed the best. However, we found that given an imperfect test, there was a trade-off between sensitivity and specificity depending on the transmission setting. In low to medium transmission settings, having an assay with a high specificity (percentage of seronegative individuals correctly identified as negative) was more important than sensitivity (percentage of seropositive individuals correctly identified as positive) for determining the vaccination recommendation (Figure 5a and b). Conversely in high transmission settings (Figure 5b and c) having an assay with a high sensitivity was more important than specificity. This result is also reflected in Figure 6 which shows the proportion of fits making the correct recommendation regarding vaccination at low (P9=30%) and high (P9=80%) transmission settings respectively. Vaccination could not be definitively excluded in low transmission settings when using a low specificity assay (*S_p_*=0.6). In high transmission settings (80% seroprevalence in 9 year olds) the correct recommendation would be to vaccinate the population. However, the opposite recommendation would be made if using a low sensitivity assay (*S_e_*=0.6) with almost all the fits recommending against vaccination. Therefore, in high transmission settings, having an assay with a high sensitivity was more important than having high specificity. Of the commercially available assays, as expected the assay with the highest sensitivity and specificity – the IgG ELISA from Standard Diagnostics Inc. (*S_e_*=98.8%, *S_p_*=99.2%), performed the best.

**Figure 5:**
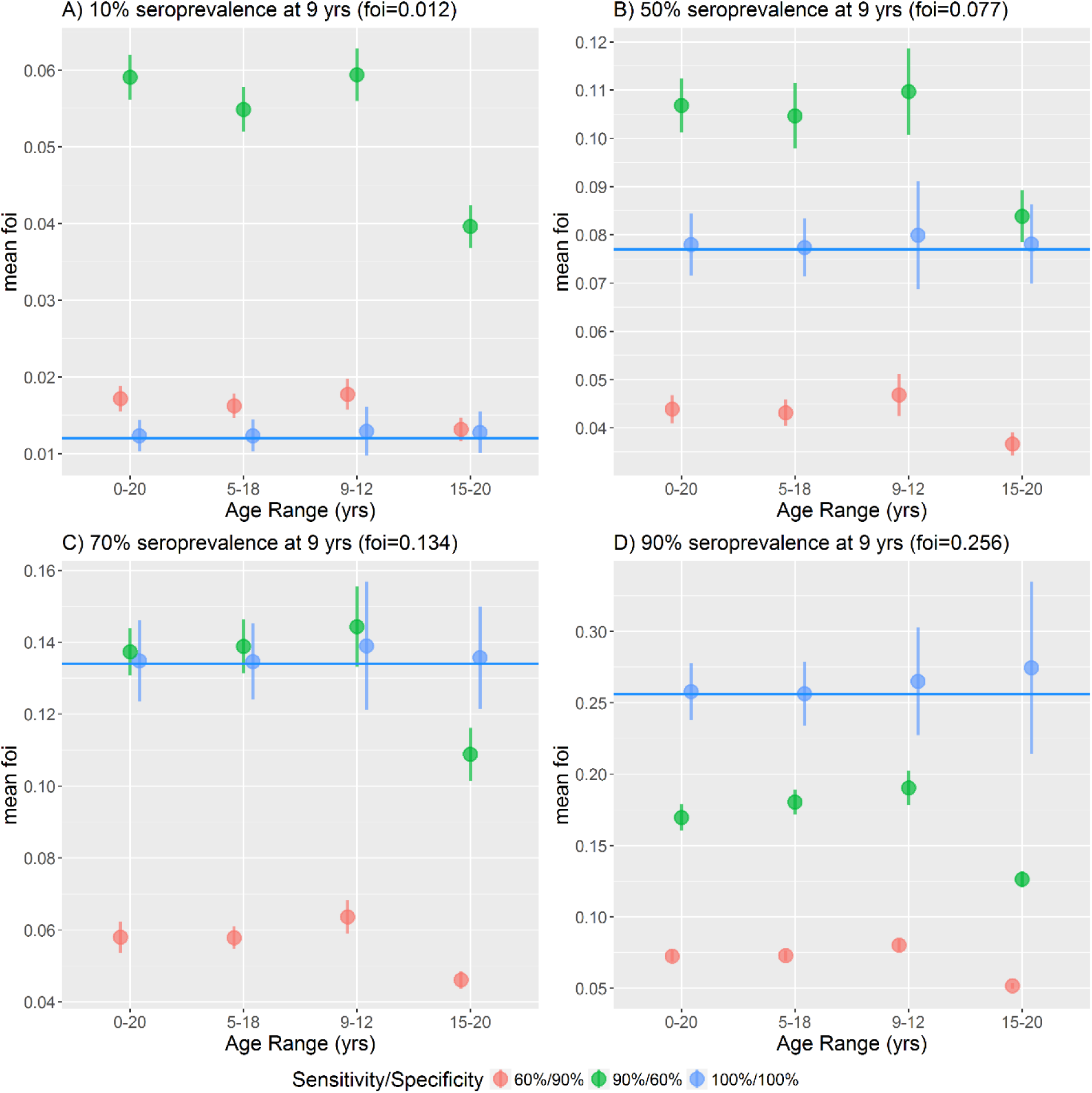
Dengue force of infection re-estimated at different transmission intensities, from a range of ages and test sensitivities and specificities. A) Very low transmission 10% seroprevalence at age 9, B) medium transmission 50% seroprevalence at age 9, C) high transmission 70% seroprevalence at age 9, D) very high transmission 90% seroprevalence at age 9. The point shows the mean of the 100 mean posterior distribution of the force of infection, the bar the standard deviation, and the horizontal blue line shows the true force of infection.

**Figure 6:**
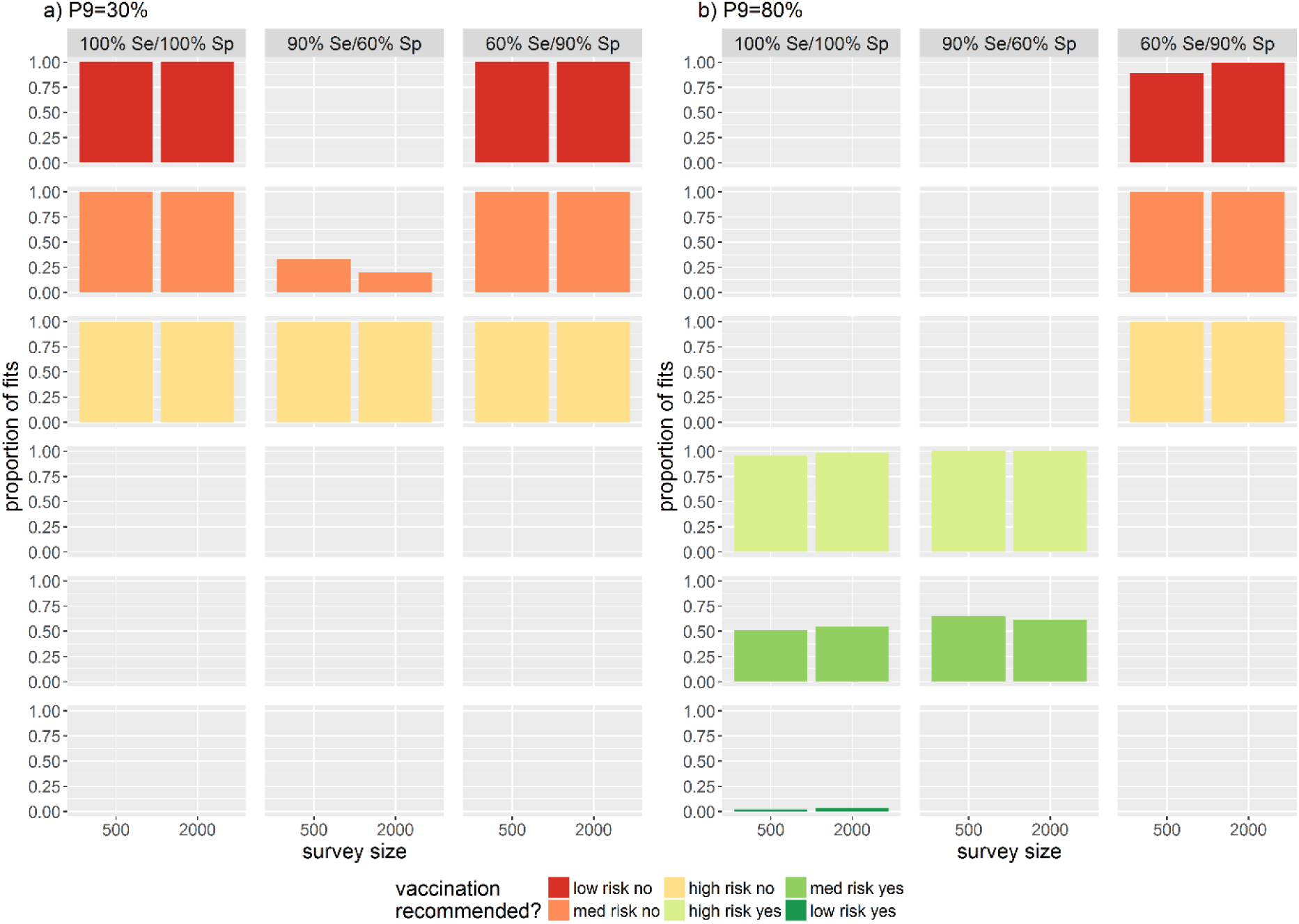
Proportion of model fits making the correct recommendation regarding vaccination when the true seroprevalence in 9 year olds is a) 30% and b) 80%, for different test sensitivities and specificities. Se = sensitivity, Sp = specificity. Red = vaccination is not recommended, green = vaccination is recommended. Low risk no = upper 95% CrI < 50%, med risk no = central estimate <50%, high risk no = upper 95% CrI < 50%, high risk yes = upper 95% CrI above 70%, med risk yes = central estimate >70%, low risk yes = lower 95% CrI >70%. CrI = credible interval.

## Discussion

The results from these simulations highlight several key considerations in the design of serosurveys to assess dengue vaccine suitability. We found that having a wide age range from which to survey was more important in accurately assessing the force of infection than a large survey size (Figure 3), with no substantial gain in precision above sample sizes of 1000 individuals. The optimum age from which to sample was dependent on the transmission setting. In high and very high transmission settings (SP9>70%) younger children under 9 years of age also need to be sampled. While in low transmission settings (SP9<30%) older aged children and even adults need to be sampled to accurately estimate the transmission intensity. The optimal test sensitivity and specificity assuming absence of a perfect test was also dependent on transmission setting. In low transmission settings, a test with a high specificity was favoured over sensitivity (Figure 5a and b). Conversely in high transmission settings, a test with high sensitivity performed better than high specificity (Figure 5b and c). Test sensitivity and specificity had a substantial impact on potential decisions about whether or not to introduce dengue vaccination in different transmission settings.

CYD-TDV is licensed for use in individuals aged 9-45 years or 9-60 years depending on the country. Our finding that it is important to sample from children younger particularly in high transmission settings is important as children <9 years old are not eligible to receive the vaccine and so are not an intuitive target population. It may also be difficult to obtain samples from this younger age group for the same reason. Conversely, in very low transmission settings, even with the widest age range of 0-20 years the transmission intensity was slightly overestimated. This suggests that older adults, in some cases above 45 years of age may also need to be sampled. This may be the case in countries such as Singapore where the seroprevalence in school-aged children is low [18–20]. In Singapore CYD-TDV has been licensed for private use only (12 years and above) with guidelines suggesting patients get pre-tested to determine past infection status [21]. However, in countries with high transmission intensity where the vaccine is intended to be used, school-based sampling should be sufficient. Ideally with younger age classes 5 years and above also included in the sampling framework [22]. Our simulations show that the 5-18 year old age group performed consistently well and this would make conducting serosurveys more feasible.

If CYD-TDV is to be introduced as a public programme, it is likely that vaccination will be rolled out at the sub-national or state level. However, dengue is geographically heterogenous even at fine spatial scales [23,24] and transmission intensity can differ substantially between sub-districts of the same administrative region [25]. Therefore, it would be beneficial to conduct multiple smaller surveys (of sufficient size) to capture the geographic heterogeneity within a region rather than to conduct one large survey in e.g. the state capital.

Our finding that optimal test sensitivity and specificity differed by transmission setting and could potentially result in false vaccination recommendations are based on two rather extreme scenarios: Se=90%/Sp=60% and Se=60%/Sp=90%. Generally, commercially available IgG ELISAs have good sensitivity and specificity and re-estimated the true transmission intensity well (Table 1). However, it should be noted that IgG ELISAs are primarily manufactured and calibrated for diagnostic purposes rather than for mass serosurveys. Additionally, anti-dengue antibodies are cross-reactive with other flaviviruses including Yellow Fever (YF), Japanese encephalitis (JE), West Nile Virus, and Zika [26–28].

Therefore, care must be taken when interpreting the results of serosurveys in areas with co-circulation of other flaviviruses, or routine JE or YF vaccination. With the emergence of Zika in South America and the large epidemic observed in 2015-2016, cross-reactivity may be of particular concern since there are currently no commercially available diagnostics or internationally recognised standards for Zika diagnosis [29]. In the context of conducting serosurveys to inform dengue vaccination, WHO recommend that a subset of samples is re-tested using neutralisation tests such as plaque-reduction neutralisation tests (PRNTs) as they do not detect the same cross-reactivity. Such validations will allow IgG ELISA cut-offs to be calibrated to the local epidemiological context. Additionally, having such baseline measurements may be important when assessing vaccine efficacy at a later date.

Our simple model assumes a constant force of infection and it is not possible to assess time or age-dependent changes in transmission intensity from single cross-sectional surveys. Dengue transmission can vary year-to-year due to the cyclical nature of transmission [30]. However, estimates derived from cross-sectional IgG surveys will still give a good estimate of the average transmission intensity over time within a population [13,31,32] which should be sufficient to assess vaccination suitability particularly in dengue endemic areas for which the vaccine is intended. Additionally, although there are known serotype-specific differences in transmission intensity non-serotype specific data can yield an estimate of the total force of infection across all serotypes consistent with the sum of serotype-specific forces of infection which can be derived from PRNT data [13].

Our main results did not change whether or not we took overdispersion into account (beta-binomial vs. binomial). However, the trends were not as marked when using a binomial model with no overdispersion (SI Text). We found that assuming greater overdispersion makes it increasingly difficult to re-estimate the transmission intensity accurately (Figure S7). The degree of overdispersion is likely to differ by setting.

Since optimal sampling age range and diagnostic sensitivity and specificity differed by transmission setting, having a method to infer likely seroprevalence using readily available data such as notification data will be important in informing serosurvey design. Previous work has shown that the dengue transmission intensity can be inferred from the age-distribution of dengue cases [33]. However, additional age-stratified seroprevalence and incidence data which are matched in time and space are required to fully validate this approach of using surveillance data as a proxy for case data.

WHO recommend that countries only consider introduction of CYD-TDV in areas “where epidemiological data indicate a high burden of disease” [34]. Although current guidance specify that the primary objective of serosurveys is to estimate the age-specific seroprevalence of dengue antibodies in a potential population, age-stratified seroprevalence surveys can show considerable fluctuations with age due to the epidemic nature of dengue transmission as well sampling variability [35]. Here we have shown that a more accurate measure of dengue endemicity is the transmission intensity or force of infection (λ) which can be estimated from such serosurveys. This average transmission intensity takes into account information from a much wider age group than a narrow vaccine target population and can then be used to better estimate the expected seroprevalence at any given age. Such data would provide better baseline estimates against which dengue surveillance systems and notification data could be verified.

Although these results are presented in the context of the CYD-TDV dengue vaccine, phase II interim analysis of the Takeda TDV vaccine has also shown lower geometric mean titres (GMT) of neutralising dengue antibodies to DENV-1 to -4 in those seronegative at baseline compared to seropositive at baseline [36]. Therefore, it is possible that similar recommendations for targeting vaccination may be made for future dengue vaccines. Finally, the considerations for serosurvey design highlighted here can be applied not just to dengue but more broadly to seroprevalence surveys in general.

